# Bacterial predation on T4 phages

**DOI:** 10.1101/2021.06.01.446436

**Authors:** Jean-Jacques Godon, Ariane Bize, Hoang Ngo, Laurent Cauquil, Mathieu Almeida, Marie Agnès Petit, Olivier Zemb

## Abstract

**Background:** Bacterial consumption of viruses has never yet been reported, even though bacteria feed on almost anything. Viruses are omnipresent predators for all organisms, but have no acknowledged active biocontrol. The viral biomass undoubtedly reintegrates the trophic cycles, however the mechanisms of this phase still remain unknown.

**Methods:** Here, we used stable isotope probing with ^13^C labelled T4 phages to monitor the increase of density of the bacterial DNA concomitant with the decrease of plaque forming units. We used ^12^C T4 phages as control.

**Results:** T4 phage disappearance in wastewater sludge was found to occur mainly through predation by *Aeromonadacea*. Phage consumption also favours significant *in situ* bacterial growth. Furthermore, an isolated strain of *Aeromonas* was observed to grow on T4 phages as sole source of carbon, nitrogen and phosphorus.

**Conclusions:** bacterial species are capable of consuming bacteriophages *in situ*, which is likely a widespread and underestimated type of biocontrol.This assay is anticipated as a starting point for harnessing the bacterial potential in limiting the diffusion of harmful viruses within environments such as gut or water.

## Introduction

For any type of bacteria, the presence of viruses may present a significant opportunity for feeding. Indeed, viruses represent 0.2 gigatons of carbon on Earth(Bar-On *et al*., 2018). For example, the major capsid protein of the T4-like bacteriophage family is one of the most prevalent proteins in the biosphere(Comeau and Krisch, 2008). Therefore phages represent a major potential carbon source into which bacteria may tap. Furthermore, viruses are also a potential source of phosphorus(Jover *et al*., 2014).

No bacterium preying on viruses have been described even though bacterial extracellular proteases are able to degrade certain bacteriophages in anaerobic wastewater treatment plants, in pure cultures(Mondal *et al*., 2015) and in soil(Nasser *et al*., 2002). In seawater, the only reported biotic pressure arises from marine ciliates that have been co-incubated with viruses and bacteria(Gonzalez and Suttle, 1993). This observation is also supported by the recent discovery of viral DNA in free-living eukaryotic cells(Brown *et al*., 2020).

Here, we show that specific bacteria can indeed degrade T4 bacteriophages *in situ*, and we confirm this observation in pure culture.

## Results

When searching for bacteriophage consumption activity, sludge from wastewater treatment plants is a relevant microbiota to investigate, as it boasts a high degradation capacity. In this work, the stable isotope probing method was applied by adding 2.2 10^10 13^C-labelled T4 phages to 200 µl of sludge corresponding to 10^8^ bacteria cells. The increase in density of the bacterial DNA of the ^12^C control bottle was then measured after the enumerated T4 phages decreased by 99%.

T4 phages were assimilated by bacteria. About 41% of the ^13^C atoms initially present in T4 phages were accounted for in the bacterial biomass. However, only nine out of the 4046 microbial species - or more accurately Amplicon Sequence Variant (ASVs) - were labelled by the ^13^C initially contained in the T4 phages (Fig. 1), thus suggesting that the incorporation of T4 phage is not a widespread ability. This incorporation generated growth, since the total biomass increased 2 fold after 24h concomitantly with the disappearance of the T4 phages.

**Figure 1.**
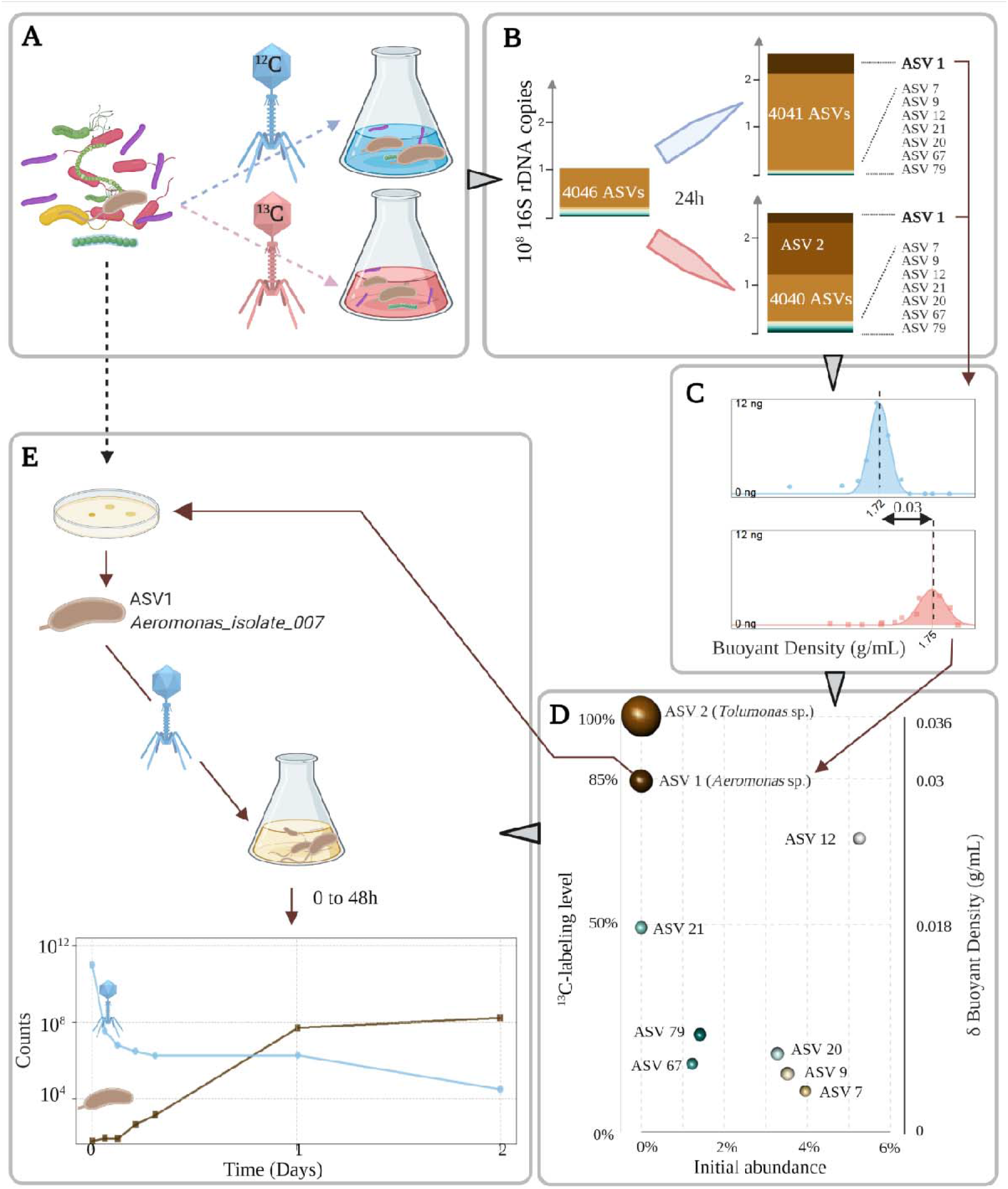
Bacterial growth on T4 phages. **A; Identification of ^13^C-labeled bacteria:** the ^13^C-labeled T4 (red) were incubated with a microbial community of wastewater treatment plant in the same conditions as the ^12^C control (blue). **B; Bacteria present in each sample:** the barplots shows the growth of each ASVs based on 16S rDNA copies, detailing the nine bacteria assimilating T4 phages. **C; The density plots show the shift in density for ASV1 (*Aeromonas* sp.)**, which was abundant in both bottles after 24h. **D;** ^**13**^**C labelling level after 24h:** the ^13^C-labeling level (left Y-axis), computed from the individual density shifts (right Y-axis), is reported against the initial abundance of the ASVs (X-axis). The volume of the spheres represents the abundance of each ASV in the ^13^C bottle. **E; *Aeromonas* sp. growth on T4 phages:** Aeromonas_isolate_007 grew on T4 phages as sole carbon and nitrogen source. When a few *Aeromonas* cells were incubated with 10^11^ T4 phages, the colony forming units (brown) increased while the plaque-forming units (blue) decreased.

The two main degraders of T4 phages were ASV1 (*Aeromonas sp*.) and ASV2 (*Tolumonas* sp.), which accounted for 5% and 29% of ^13^C atoms found in the bacterial biomass respectively. Both belong to the *Aeromonadaceae* family and exhibit strong growth rates. Indeed, both rose from undetectable levels to 51% of the biomass, while the density of their DNA increased because they incorporated ^13^C atoms from the isotopically labelled T4 phages. For example, the 2 10^6^ *Aeromonas* cells present after 24h contained 85% of ^13^C atoms in their DNA whose density shifted from 1.72 g/mL to 1.75 g/mL in the bottle with ^13^C-labeled T4 phages. The 16S rRNA sequences assigned to *Aeromonas* represented 19% and 8% of the total reads in the ^12^C and the^13^C bottles respectively, thus revealing a consistent growth from initially undetectable levels.

In addition to the *Aeromonadaceae* family, two species (ASV12 and 21) belonging to the Ignavibacteriales PHOS-HE36 family, although labelled with medium strength (49 and 71%) and negligible growth (0 and 5.87 10^6^ synthetized cells respectively), still gathered 5% of the ^13^C atoms. The last 5 species with significant DNA density shifts (ASV7, 9, 20, 67 and 79) accounted for the remaining 1% of ^13^C atoms but their weak labelling level may have resulted from indirect labelling.

To confirm the quality of *Aeromonas sp*. as a predator of T4 phages, an *Aeromonas*-selective medium was used for retrieving an *Aeromonas* colony from the initial sludge and called it Aeromonas_isolate_007. The analysis of the whole genome confirmed that this isolate belongs to an intermediate clade between *Aeromonas media* and *Aeromonas rivipollensis* species. Aeromonas_isolate_007 was incubated with T4 phages as only substrate. Starting with 50 resting bacterial cells, the population reached 1.6 10^8^ cells after 24 hours at 20°C while consuming 10^11^ T4 phages (Fig. 1C). No growth was observed when T4 phages were absent.

*Aeromonas sp*. can also capture T4 phages when their concentrations were comparable with environmental conditions: 7 10^4^ T4 phages/mL decreased to 2 10^3^ T4 phages/mL when incubated with Aeromonas_isolate_007 cells (Fig 1C). No decrease in the T7 phage has been observed in similar experiments where the T4 phage was replaced by the T7 phage.

## Discussion

*Aeromonas* cells are present in virtually any environment(Janda and Abbott, 2010), including wastewater treatment plants where their abundance is around 0.1 %(Ye *et al*., 2012).

Interestingly, *Aeromonas* cells have an S-layer(Noonan and Trust, 1997) associated with lipopolysaccharides (Sleytr *et al*., 2014) and an outer membrane protein C, which are known to bind the T4 phages to the surface of *E*.*coli* cells(Islam *et al*., 2019). Once captured at the surface, the phage is likely degraded by several extracellular enzymes, including DNase and protease (Janda, 1985). For example, metallo- and serine-proteases found in *Aeromonas* are involved in the degradation of large molecules such as albumin, earning the nickname of ‘Jack-of-all-trades’ due to this enzymatic versatility(Seshadri *et al*., 2006). Finally, *Aeromonas* possesses transporters to uptake the resulting amino acids and peptides (Seshadri *et al*., 2006).

Bacterial predation on bacteriophages is rich in consequences because bacteriophages are ultimate predators at the top of all food chains since they are not hunted. Indeed bacteriophage decay is mainly considered abiotic via adhesion to particulate material, chemical inactivation or degradation by solar radiation or passive grazing by flagellates(González and Suttle, 1993). In the oceans, this predation likely allows for the upper levels of the trophic chain carbon to access to the 7% of dissolved nitrogen, the 5% of phosphorus and the 1% of dissolved organic carbon contained in the viral particles (Jover *et al*., 2014).

Furthermore, the diversity in bacteriophages could be partly related to the presence of phage-specific bacterial bacteriophage-hunters. Indeed, the bacterial predators of T4 phages do not appear to consume T7 bacteriophages. Therefore, brutal increase of a specific phage in the environment could be specifically controlled by a phage-eating bacterium, forming a killing-the-killer loop.

In conclusion, bacteria that are capable of eliminating specific viruses changes our vision of the food webs and represent a noteworthy avenue to explore to control harmful viruses such COVID-19, bacteriophages that disrupt dairy fermentations or rotaviruses causing diarrhoea.

## Supporting information

suppelmental table

## Declarations

### Ethics approval and consent to participate

Not Applicable

### Consent for publication

Not Applicable

### Availability of data and materials

High-throughput sequencing data have been deposited on NCBI (https://www.ncbi.nlm.nih.gov/bioproject) under accession number PRJNA650397 and the genome of Aeromonas_isolate_007 is accessible with the BioSample accession number SAMN17689348.

### Contributions

J.J.G., O.Z., A.B. and N.H. designed and performed the experiments; J.J.G., O.Z., L.C. and M.A. performed computational experiments; J.J.G., O.Z., L.C., M.A. and M.A.P. contributed lineaging data and expertise; J.J.G. and O.Z. prepared the manuscript with assistance from all authors; J.J.G. and O.Z. supervised and directed the research.

## Acknowledgments

We thank Caroline Achard for insights on bacteriophage capture, Martin Beaumont and Amira Bousleh for the qPCR of the internal standard. Electronic microscopy work has benefited from the facilities and expertise of MIMA2 MET – GABI, INRA, Agroparistech, 78352 Jouy-en-Josas, France. Sequencing was performed in collaboration with the GeT core facility, Toulouse, France (http://get.genotoul.fr) supported by France Génomique National infrastructure, funded as part of “Investissement d’avenir” program managed by the Agence Nationale pour la Recherche (contract ANR-10-INBS-09). Figure 1 was created by Biorender.

## Authors’ information

JJG directed the PhD of OZ in bacterial ecology in France. In a following postdoc, OZ learned stable-isotope probing in sludge in Australia and was granted a small project to study the adsorption of T2 phages in the light of electrostatics. The combination of SIP and phage expertise turned out useful to answer JJG’s question about the predation of phages together with AB, which has expertise in producing purified isotopically-labeled phages and MAP, which could confirm the results in pure culture with JJG since an isolate was successfully isolated from the sample. MA’s expertise in genome annotation was put to use to speculate about the mechanisms involved in the capture, digestion and assimilation of T4 phages by bacteria.

