## Supplementary material for "Bacterial predation on T4 phages": suppelmental table

**Supplementary information**

TABLE OF CONTENT

### 1 ****Stable isotope probing experiment on the initial sample****

#### 1.1 **Preparation of the ^13^C-labeled T4 bacteriophages**

T4-phage particles labeled with ^13^C were produced on *Escherichia coli* B cells (DSM 613) grown in M9 minimal medium with ^13^C-glucose as the sole carbon source. The M9 medium was prepared using M9, Minimal Salts, 5X (Sigma-Aldrich), by adding MgSO4 (Sigma-Aldrich) and CaCl2 (Sigma-Aldrich) at final concentrations of 1 mM, and D-Glucose at a final concentration of 10 g/L. Moreover, additional salts were added to favor phage adsorption (solution of CaCl2 0.5 M and MgCl2 1M, diluted 1 000 times in the culture medium). More precisely, starting from an *E. coli* stock of cells frozen in LB and glycerol, two successive overnight pre-cultures were grown in LB medium (LB broth, Fisher). Subsequently, 5 x 20 mL of M9 minimal medium containing D-Glucose-^13^C6 as the sole carbon source (10 g/L) were each inoculated with 20 µL of the second *E. coli* pre-culture; approximately 1500 T4-phage particles (DSM 4505, in PFU) were added. Finally, 20 µL of a solution containing 0.5 M CaCl2 and 1M MgCl2 were also added in each case to favor phage adsorption.

After 30 hours of incubation at 37°C under agitation, the T4 phage particles were collected: the cultures were centrifuged during 15 minutes at 5 000 g, 10°C. The supernatants were collected and filtered at a 0.22 µm pore-size (PES filters, Milipore). They were subsequently incubated overnight in 8% w/v PEG 6000 and 0.5 M NaCl solution, at 4°C, to precipitate viral particles. The supernatants were centrifuged at 20 000g, during 30 minutes, at 4°C. The pellets were suspended in SM buffer (100 mM NaCl, 8 mM MgSO4, 50 mM Tris pH 7.5), and centrifuged once more at 20 000 g for 4h at 4°C. Viral particles were finally suspended in 1.4 mL of SM buffer and stored at 4°C before use.

To obtain unlabeled T4 phage particles, the same procedure was used except that unlabeled glucose was employed in the M9 minimal medium.

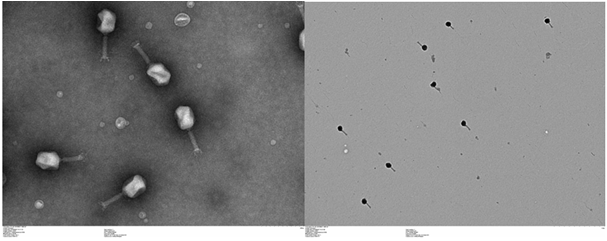

Figure S 1 : Purity of the bacteriophage preparation

*We checked the purity of the bacteriophage preparation by electronic microscopy. Materials were directly adsorbed onto a carbon film membrane on a 300-mesh copper grid, stained with 1% uranyl acetate, dissolved in distilled water, and dried at room temperature. Grids were examined with Hitachi HT7700 electron microscope operated at 80kV (Elexience – France), and images were acquired with a charge-coupled device camera (AMT).*

Table S 1 : Decrease of the free ^13^C and ^12^C bacteriophages by PFU of the supernatant. We indicate the absolute numbers of phages and their percentages compared to t_0_.

| Time (min) | T4 phages  in ^12^C bottle | T4 phages  in ^13^C bottle | T4 phages  in ^12^C bottle | T4 phages  in ^13^C bottle |
| --- | --- | --- | --- | --- |
| 0 | 2.8 10^10^ | 2.2 10^10^ | 100 % | 100 % |
| 24 | 5.2 10^8^ | 9.4 10^8^ | 1.9 % | 4.2 % |
| 122 | 2.6 10^7^ | 4.1 10^8^ | 0.9 % | 1.8 % |
| 445 | 3.2 10^7^ | 3.3 10^8^ | 1.2 % | 1.5 % |
| 1375 | 5 10^2^ | 5.7 10^5^ | 0 % | 0 % |

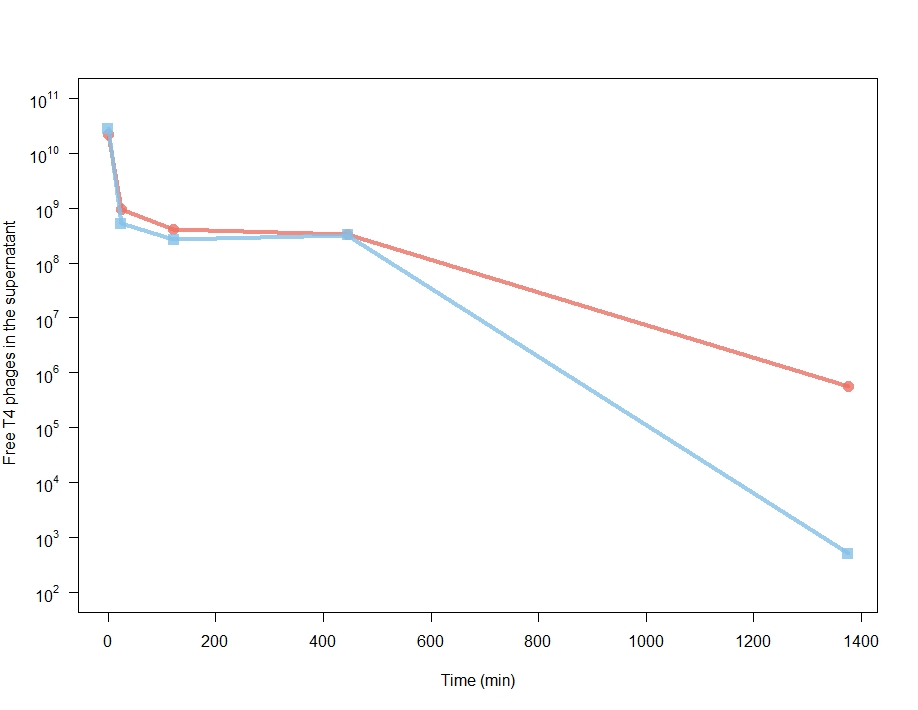

Figure S 2: Decrease of the free ^13^C and ^12^C bacteriophages by PFU of the supernatant

*Free purified T4 phages were incubated with the initial sample of wastewater treatment plant. The initial concentration was 2.24 10^10^ T4 phages in 5 mL, i.e. 4.48 10^9^ T4 phages/mL. After 24h, over 99% of the T4 phages are missing from the supernatant due to incorporation or absorption into the microbial biomass. The red curve represents the ^13^C bottle and the blue curve represents the ^12^C bottle.* *The PFU titers of the obtained 13C- and unlabeled T4 stock solutions were determined on a bacterial lawn of E. coli cells (DSM 613), using the soft-agar overlay technique. More precisely, 5 µL of T4 phage stock solutions and serial dilutions of them (factor 10 or 15) were plated a soft layer containing 7.5 g/L of Agar and E.coli cells (DSM 613) pre-cultured in LB medium, in LB-Agar plates (15 g/L of Agar, Sigma-Aldrich). After a short period of drying, the Petri dishes were incubated at 37°C during 24 hours in static conditions. The PFU titers were determined by counting the visible plaques and calculating the concentration in the original stock solutions.*

#### 1.2 **Detailed Calculations of the ^13^C mass balance**

This paragraph explains in details the calculations reported in Table S3. For a simpler explanation, we will focus here on the calculations about ASV1 (*Aeromonas* sp.).

**STEP1:** using qPCR, we determine the absolute abundance of bacteria in the ^13^C bottle at the beginning of the experiment and at the end of the experiment. This can be done because we spiked 3 10^5^ copies of a synthetic DNA standard to 200 µl of the initial sample ^1^, which we extracted immediately and quantified using 16S universal primers. We also spiked the final pellet with 3 10^5^ copies of a synthetic DNA standard and quantified it using specific primers. The comparison of the 16S and internal standard qPCR curves indicates that the initial sample contained 1.798^23.462^/1.828^13.215^=326 fold more bacterial 16S rDNA copies than internal standard (Table S 2), so that the tube contained 3 10^5^ x 326 = 9.78 10^7^ copies of bacterial 16S at the beginning of the experiment. At the end of the experiment, comparing the 16S and internal standard qPCR curves indicates that bacterial 16S rDNA were 783 fold more abundant than the spiked synthetic standard, leading to an estimated 2.35 10^8^ copies of bacterial 16S rDNA genes in the ^13^C bottle. For an accurate mass balance, we also consider the fact that 240 µl were taken out for PFU measurements during the experiment, so that we estimate that 2.47 10^8^ copies of 16S rDNA would have been present at the end of the experiment in the ^13^C bottle if no sampling had been performed. Since we used the LinReg software, we also accounted for the slight individual variations in qPCR efficiency, which were between 80 and 84% for the internal standard and between 80 and 82% for the bacterial 16S rRNA genes respectively. In total, we estimate a global biomass increase of 2.35 fold. This global increases regroups ASVs that are initially abundant and grow slightly with ASVs that are initially rare and grow massively.

Table S 2 : Quantification of the bacterial density with the internal standard at the beginning and at the end of the experiment.

| Sample name | Primers | Efficiency | Cycle Threshold | Spiked internal standard | 16S rDNA |
| --- | --- | --- | --- | --- | --- |
| Initial_sample | Internal standard | 79.8 % | 23.462 | 3 10^5^ copies | 9.78 10^7^ copies |
|  | V4V5 | 82.8% | 13.215 |  |  |
| Final_^13^C_sample | Internal standard | 80.1% | 14.133 | 3 10^5^ copies | 2.35 10^8^ copies |
|  | V4V5 | 84.5% | 24.461 |  |  |

**STEP2:** using 16S metabarcoding, we use the absolute abundance of 16S rDNA of each ASV at the beginning and at the end of the experiment to estimate the number of 16S rDNA copies produced during the experiment for each ASV. ASV1 (*Aeromonas* sp.) was undetectable in the 3748 sequences of 16S rRNA genes obtained at the beginning of the experiment. At the end, 303 sequences out of 3578 (i.e. 8%) belonged ASV1 (*Aeromonas* sp.). For the two instances where the final 16SrDNA abundances were lower than the initial 16S rDNA abundances (possibly due to part of the population dying combined with another part showing a small growth), we neglected the contribution of these ASVs to the ^13^C mass balance.

**STEP3:** using a database of 16S rDNA copy number^2^, we convert the increase in 16S rDNA into the number of cells. *Aeromonas* has 10 copies of 16S rDNA, so that 2.09 10^7^ copies of 16S rDNA corresponds to 2.09 10^6^ *Aeromonas* cells. ASV 1 was undetectable at t_0_, so that if only one cell of ASV 1 (*Aeromonas* sp.) was present at the beginning of the experiment, 22 generations would have been needed in the course of the experiment to produce 2 10^6^ cells, i.e. 62 minutes per generation. If ASV 1 (*Aeromonas* sp.) was just below the detection limit ((1/3748*9.78 10^7^)/10= 2600), corresponding to an average growth rate of 125 minutes per generation.

**STEP4:** Assuming that each cell contains 30 fg of carbon, we convert the number of cells produced during the course of the experiment to a total carbon reservoir at the end of the experiment (^13^C + ^12^C). *Aeromonas* has 2.09 10^7^ cells at the end, and a negligible amount at the beginning (undetectable). We then estimate the amount of total carbon that was captured by the growth of ASV1 (*Aeromonas* sp.). For example, 2.09 10^7^ *Aeromonas* cells translate into 6.28 10^-8^ g of total carbon content. It should be noted that 30 fg of carbon was measured for dried *Aeromonas* cells using a Leco CHN analyzer ^3^ but we assume the same value for every ASV. In the aforementioned work, the cellular carbon content varied between 20 and 40 fg, depending on the bacterial species.

**STEP5:** using the shifts in buoyant density between the ^12^C- and the ^13^C- bottles, we estimate the labeling level of each ASV so that we can estimate their contribution to the ^13^C mass balance. We compare the buoyant density of each ASV in the ^13^C bottle to the density of in the ^12^C bottle by fitting a normal curve to the absolute numbers of 16S rRNA genes that were detected in each fraction. Since we measured the density of each fraction by refractometry, the mean of the normal curve is the best estimate of the actual buoyant density of the ASV (Figure S 3). For example, the 16S DNA of ASV1 (*Aeromonas* sp.) have a mean density of 1.72 in the ^12^C bottle and 1.75 in the ^13^C bottle with good fits (R²=0.98 and 0.89). As mentioned in the text, *ASV2 (Tolumonas* sp.) is not abundant enough in the ^12^C bottle to fit a normal curve on its distribution across the gradient, so we used its theoretical density based on its GC content (which shows a 15% error). The shift between the ^12^C- and ^13^C- densities was converted into a percentage of ^13^C by dividing by 0.036 ^4^. The labeling level of ASV1 (*Aeromonas* sp.) was (1.75-1.72)/0.036=85%.

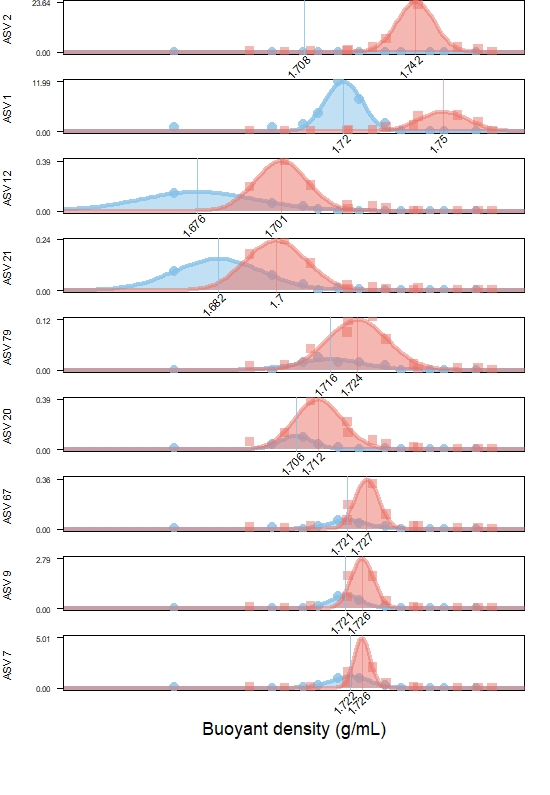

Figure S 3 : Gaussian fits of the 9 most labeled ASVs.

*The amount of DNA for the top 9 ASVs (in ng) is plotted against the density of the Cesium gradient (in g/mL). The blue color is in the control bottle. The red color indicates the amount of DNA in the bottle supplemented with 3.2 µg of 13C in the 13C- labeled T4 bacteriophages. The dots indicate the actual measures performed after 24h, i.e. the amount of DNA of each ASV (Qbit was combined with the 16S rDNA sequencing) in each fraction whose density was measured by refractometry. The lines indicate the Gaussian distributions to accurately estimate the mean buoyant density. The mean buoyant density of ASV 2 (Tolumonas sp.) was estimated with the theoretical value in the 12C bottle, as ASV2 did not grow sufficiently in that bottle to fit a reliable Gaussian fit. The pie charts indicate the ratios of the nine 13C-labeled ASV in the 13C and 12C bottles after 24h and their proportion in the initial sample is reported in the X axis. For example, the pie of Aeromonas is roughly balanced because ASV 1 (Aeromonas sp.) represented 19% and 8% in the 12C and 13C bottles respectively. In contrast, ASV 2 (Tolumonas sp.) only grew substantially in the 13C bottle (red).*

**STEP6:** To complete the mass balance of the ^13^C-atoms of the labeled T4 phages, we multiply the total carbon content of each species by its labeling level. For example, ASV1 (*Aeromonas* sp.) represents 6.28 10^-8^ x 0.85 = 5.33 10^-8^ g of carbon (Table S3).

**STEP7:** To estimate the mass of carbon needed to account for the carbon content observed each species, we assume that the carbon use efficiency for each ASV is 33% (see pure culture experiment). It should be noted that we measured the yield for ASV1 (*Aeromonas* sp.) and we then we assume the same yield for every ASV. Once corrected by the carbon use efficiency (i.e. the bacterial yield), the estimation of ^13^C needed by each species is compared to the 3.2 10^-6^ g of ^13^C incorporated in the 2.28 10^10^ ^13^C-labeled T4 phages (since each T4 viral particle contains 1.49 10^-16^ g C and we assume that they were 100% labeled with ^13^C because of their production method). For example, we estimated 2.09 10^6^ newly synthesized cells of ASV1 (*Aeromonas* sp.), which account for 6.28 10^-8^ g ^13^C therefore correspond to 5% of the ^13^C atoms initially present. Adding the contributions of the 9 most-labeled ASVs accounts for 41% of the initial mass of ^13^C (Table below).

Table S 3 : ^13^C mass balance of the isotopically labeled T4 bacteriophages.

*This table indicates the rationale for the ^13^C mass balance following the steps described above. For example, ASV 1 (Aeromonas sp.) represents 8% of the 2.47 10^8^ 16S rDNA copies found at the end of the experiment, which represents 5% of the total amount of ^13^C present in the initial T4 phages because the 30fgC-cells labeled at 85% needed 5.3 10^-8^g of ^13^C if we consider a 33% yield.*

| STEP | Description | ASV 1 | ASV 2 | ASV 7 | ASV 9 | ASV 12 | ASV 20 | ASV 21 | ASV 67 | ASV 79 |
| --- | --- | --- | --- | --- | --- | --- | --- | --- | --- | --- |
| 1 | Absolute number of 16S rDNA copies in the initial sample | 9.78 10^7^ | | | | | | | | |
|  | Absolute number of 16S rDNA copies in the final sample | 2.47 10^8^ | | | | | | | | |
| 2 | Counts in the initial sample  (out of 3748 sequences) | 0 | 0 | 148 | 132 | 197 | 123 | 0 | 46 | 53 |
|  | Counts in the final ^13^C sample  (out of 3578 sequences) | 303 | 1545 | 37 | 62 | 52 | 55 | 55 | 41 | 55 |
|  | Relative initial abundance of 16S rDNA of each ASV | 0% | 0% | 4% | 4% | 5% | 3% | 0% | 1% | 1% |
|  | Relative final abundance of 16S rDNA of each ASV | 8% | 43% | 1% | 2% | 1% | 2% | 2% | 1% | 2% |
|  | Absolute number of 16S rDNA copies in the initial sample of each ASV | 0 | 0 | 3.86 10^6^ | 3.44 10^6^ | 5.14 10^6^ | 3.21 10^6^ | 0 | 1.20 10^6^ | 1.38 10^6^ |
|  | Absolute number of 16S rDNA copies in the final sample of each ASV | 2.09 10^7^ | 1.07 10^8^ | 2.55 10^6^ | 4.28 10^6^ | 3.59 10^6^ | 3.80 10^6^ | 3.80 10^6^ | 2.83 10^6^ | 3.80 10^6^ |
|  | Number of newly synthetized 16S copies | 2.09 10^7^ | 1.07 10^8^ | 0 | 8.36 10^5^ | 0 | 5.87 10^5^ | 3.80 10^6^ | 1.63 10^6^ | 2.41 10^6^ |
| 3 | Number of 16S rDNA copies per genome of each ASV ^2^ | 10 | 10 | 4 | 4 | 1 | 2 | 1 | 2 | 2 |
|  | Number of newly synthetized cells of each ASV | 2.09 10^6^ | 1.07 10^7^ | 0 | 2.09 10^5^ | 0 | 2.94 10^5^ | 3.80 10^6^ | 8.15 10^5^ | 1.21 10^6^ |
| 4 | Carbon content (g/cell) | 3.00 10^-14^ | | | | | | | | |
|  | Total Carbon content in each ASV (g) | 6.28 10^-8^ | 3.20 10^-7^ | 0 | 6.27 10^-9^ | 0 | 8.81 10^-9^ | 1.14 10^-7^ | 2.45 10^-8^ | 3.62 10^-8^ |
| 5 | Mean 12C density (g/mL) | 1.72 | 1.7 | 1.72 | 1.72 | 1.68 | 1.71 | 1.68 | 1.72 | 1.72 |
|  | Goodness_fit_in_^12^C (R²) | 0.98 | 0.4 | 0.99 | 0.97 | 0.99 | 0.97 | 0.98 | 0.88 | 0.89 |
|  | Mean corrected ^12^C density | 1.72 | 1.71 | 1.72 | 1.72 | 1.68 | 1.71 | 1.68 | 1.72 | 1.72 |
|  | Mean ^13^C density | 1.75 | 1.74 | 1.73 | 1.73 | 1.7 | 1.71 | 1.7 | 1.73 | 1.72 |
|  | Goodness_fit_in_^13^C (R²) | 0.89 | 0.99 | 0.9 | 0.89 | 0.98 | 0.93 | 0.99 | 0.94 | 0.94 |
|  | Labeling Level (%) | 85% | 95% | 10% | 14% | 71% | 19% | 49% | 16% | 23% |
| 6 | ^13^C carbon content in each ASV (g) | 5.3 10^-8^ | 3.0 10^-7^ | 0 | 8.8 10^-10^ | 0 | 1.7 10^-9^ | 5.6 10^-8^ | 3.9 10^-9^ | 8.3 10^-9^ |
| 7 | Carbon use efficiencicy  (i.e. Bacterial yield) | 0.33 | | | | | | | | |
|  | Contribution to the ^13^C mass balance (out of the 3.2 µg of ^13^C in the bacteriophages) | 5%^1^ | 29%^1^ | 0%^1^ | 0%^1^ | 0% | 0% | 5% | 0% | 1% |
|  | ^13^C mass balance | 41% | | | | | | | | |

^1^, 83% of the predation of T4 phages is due to *Gammaproteobacteria*, which also include *E. coli*, the natural host of T4 phages.

#### 1.3 **Taxonomy of the ^13^C-labeled ASVs**

Table S 4 : Taxonomic affiliation of the ASVs that are significantly labeled with ^13^C

*This table indicates the taxonomy of the labeled ASV performed by DADA2 with the Siva138 dataset. The taxonomy was also checked by blasting on the NCBI database.*

| seq_ID.x | Class | Order | Family | Genus |
| --- | --- | --- | --- | --- |
| ASV 2 | Gammaproteobacteria | Aeromonadales | Aeromonadaceae | Tolumonas |
| ASV 1 | Gammaproteobacteria | Aeromonadales | Aeromonadaceae | Aeromonas |
| ASV 12 | Ignavibacteria | Ignavibacteriales | PHOS-HE36 | NA |
| ASV 21 | Ignavibacteria | Ignavibacteriales | PHOS-HE36 | NA |
| ASV 79 | Bacteroidia | Chitinophagales | Saprospiraceae | Haliscomenobacter |
| ASV 20 | Bacteroidia | Chitinophagales | Saprospiraceae | NA |
| ASV 67 | Anaerolineae | Ardenticatenales | NA | NA |
| ASV 7 | Gammaproteobacteria | Burkholderiales | Rhodocyclaceae | NA |
| ASV 9 | Gammaproteobacteria | Burkholderiales | Rhodocyclaceae | Dechloromonas |

### 2 ****Isolation of Aeromonas_isolate_007 and subsequent experiments****

**Following the SIP experiment, we could luckily isolate a strain of Aeromonas sp (ASV1) from the initial sample using** the **Aeromonas Isolation Agar medium (Sigma 17118) with ampicillin since Aeromonads are resistant to ampicillin. Therefore, we could confirm that *Aeromonas* sp. was indeed able to assimilate the carbon of the T4 phages. Furthermore, we could show that *Aeromonas* could use T4 phages as carbon and nitrogen source with a 33% yield and scan the genome of Aeromonas_isolate_007 for putative mechanisms by which *Aeromonas* could capture, digest the T4 phage proteins and transfer the generated peptides into the intracellular space.**

#### 2.1 **Pure culture experiment**

We incubated 50 *Aeromonas* cells with 10^11^ T4 phages in 1mL. After 24h, we counted 1.63 10^8^ Aeromonas cells. Converting 10^11^ T4 phages into 1.6 10^8^ ASV1 (*Aeromonas* sp.) cells (Figure S 4) corresponds to 9.14 10^-14^ g of carbon (613 bacteriophages of 1.49 10^-16^ gC) per *Aeromonas* cell containing 3 10^-14^ of carbon, hence a carbon use efficiency of 33% (also called bacterial yield). The number of generations in 2880 minutes was estimated to (ln 1.6 10^8^/58)/ln(2) = 21.3, hence 134 minutes per generation.

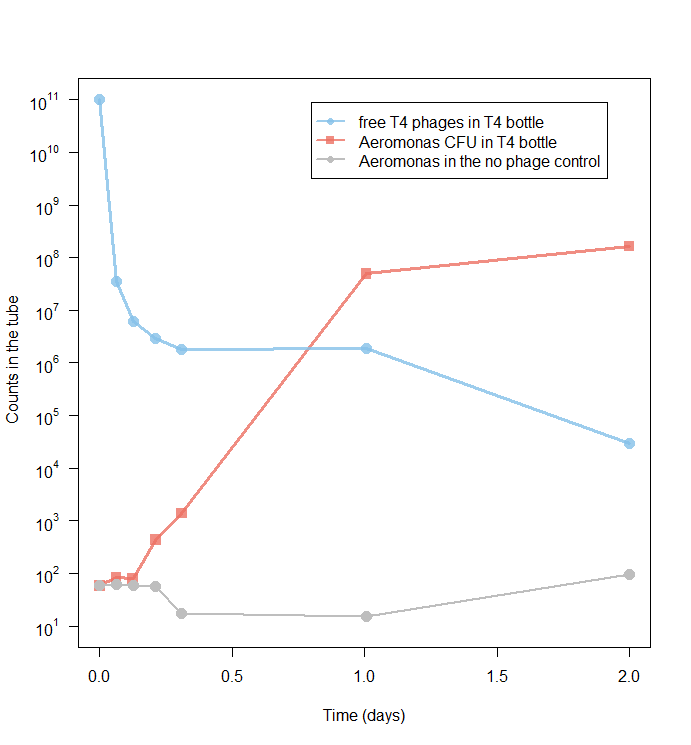

Figure S 4 : Bacteriophage decrease and Aeromonas_isolate_007 growth in pure culture

The red line indicates the number of colony-forming units over time when 50 Aeromonas cells are supplemented with 10^11^ T4 bacteriophages (blue line). The grey line demonstrates that the SM buffer in which the T4 bacteriophages are conserved is not responsible for the growth.

#### 2.2 **Whole genome sequencing**

##### 2.2.1 **Sequencing technique and statistics**

**The whole genome sequencing was performed on the GeT-Plage facility in Toulouse.**

**Briefly, Aeromonas DNA was fragmented by sonication and sequencing adaptators were ligated. 8 cycles of PCR were applied to amplify libraries. Library quality was assessed using an Advanced Analytical Fragment Analyzer and libraries were quantified by QPCR using the Kapa Library Quantification Kit. DNA-seq experiments were performed on an Illumina Miseq using a paired-end read length of 2x300 pb with the Illumina MiSeq Reagent Kits v3. The sequences were quality trimmed with fastp v0.20.0****^5^****, assembled by Spades v3.14.1****^6^** **after removing the residual phiX by using bowtie2 v2.3.5.1****^7^****, and filtering scaffolds smaller than right insert size quantile 525nt and coverage smaller than 50X. The assembly statistics of the *Aeromonas* genome are below:**

- Total genome size : 4667413 nt

- Max scaffold size : 1964260 nt

- Min scaffold size (after scaffold coverage and size filtering) : 504 nt

- Nb scaffold : 29

- Nb contig : 31

- N50 scaffold size : 947468 nt

- Average scaffold size : 160945.28 nt

- Nb Total gene : 4172

- % complete gene : 99.73

- 16S rRNA nb fragment :  2

- 16S rRNA fragment size : 1016 nt + 531 nt

- nb of 16S rRNA copies based on ratio with scaffold coverage : 10.89

- nb of 16S rRNA copies based on reference database ^2^: 10

##### 2.2.2 **Phylogenetic tree of Aeromonas_isolate_007**

**The whole genome sequencing of the Aeromonas_isolate_007 by Illumina Miseq narrows down the phylogeny of the strain and offers suggestions with respect to degradative enzymes that may help bacteriophage digestion Table S5).**

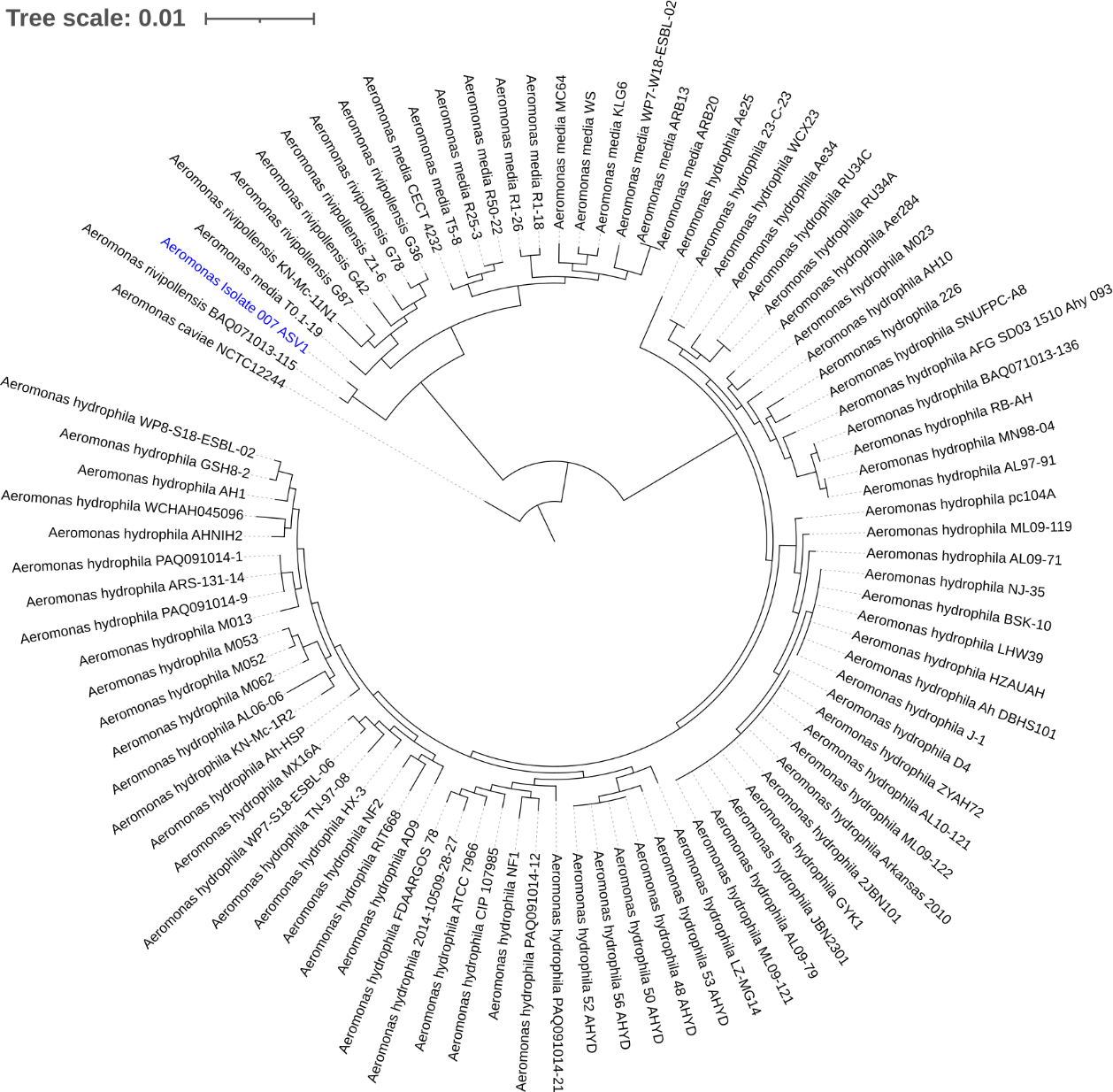

Figure S 5 : Phylogenetic tree of Aeromonas_Isolate_007_151020

*The analysis of the whole genome confirms that Aeromonas_Isolate_007_151020 belongs to an intermediate clade between Aeromonas media and Aeromonas rivipollensis species.*

##### 2.2.3 **Schematic model of the phage predation**

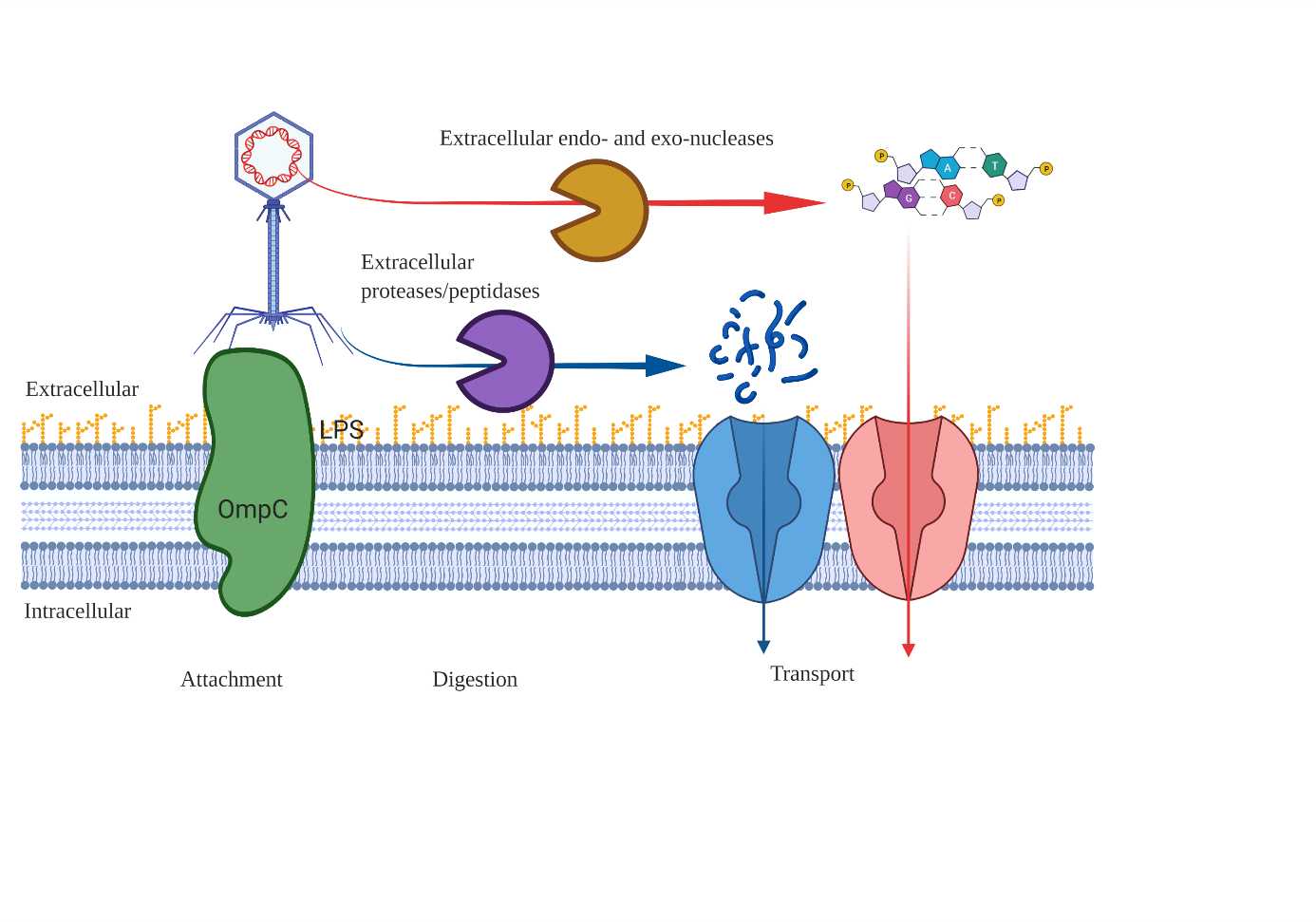

Figure S6: Schematic representation of the capture, digestion and absorption of the T4 phages by Aeromonas sp

*Since T4 phages are large particles (not to scale in the figure)., we assume that Aeromonas captures the T4 phages due to its similarity with the cell wall of E.coli. Once captured, the proteins of the capsid may be digested by the extracellular proteases present in Aeromonas sp and the DNA could be degraded as well. The products of these degradations could be ingested by Aeromonas via common transporters.*

##### 2.2.4 **Annotation of the putative genes involved in phage predation**

The functional annotation was performed with RAST8, eggNOG-mapper (v2.0.0)9 and TMHMM10 to predict the extracellular location of the proteins. The results are reported in the table below.

Table S 5 : Annotation of the Aeromonas proteins putatively involved in the capture, digestion and absorption of T4 phages, which are reported in the table below.

| **Putative function** | **RAST protein id** | **contig id** | **RAST_function** | **T**MHMM **inside** | **T**MHMM **transmembrane** | **T**MHMM **outside** |
| --- | --- | --- | --- | --- | --- | --- |
| CAPTURE | fig\|642.770.peg.2774 | NODE_2_length_947468_cov_91.192050 | Outer_membrane_porin_OmpC | 1 | 1 | 1 |
|  | fig\|642.770.peg.1967 | NODE_1_length_1964260_cov_90.024368 |  | 0 | 0 | 1 |
|  | fig\|642.770.peg.3181 | NODE_3_length_439005_cov_94.027299 |  | 0 | 0 | 1 |
| Extracellular DIGESTION of large proteins | fig\|642.770.peg.202 | NODE_1_length_1964260_cov_90.024368 | putative_extracellular_serine_protease | 0 | 0 | 1 |
|  | fig\|642.770.peg.309 | NODE_1_length_1964260_cov_90.024368 | Uncharacterized_protease_YhbU | 0 | 0 | 1 |
|  | fig\|642.770.peg.653 | NODE_1_length_1964260_cov_90.024368 | Tail-specific_protease_precursor_(EC_3.4.21.102) | 0 | 0 | 1 |
|  | fig\|642.770.peg.664 | NODE_1_length_1964260_cov_90.024368 | Lon_protease_homolog_YcbZ | 0 | 0 | 1 |
|  | fig\|642.770.peg.903 | NODE_1_length_1964260_cov_90.024368 | ATP-dependent_protease_La_(EC_3.4.21.53)_Type_I | 0 | 0 | 1 |
|  | fig\|642.770.peg.904 | NODE_1_length_1964260_cov_90.024368 | ATP-dependent_Clp_protease_ATP-binding_subunit_ClpX | 0 | 0 | 1 |
|  | fig\|642.770.peg.905 | NODE_1_length_1964260_cov_90.024368 | ATP-dependent_Clp_protease_proteolytic_subunit_ClpP_(EC_3.4.21.92) | 0 | 0 | 1 |
|  | fig\|642.770.peg.919 | NODE_1_length_1964260_cov_90.024368 | Protease_III_precursor_(EC_3.4.24.55) | 0 | 0 | 1 |
|  | fig\|642.770.peg.976 | NODE_1_length_1964260_cov_90.024368 | ATP-dependent_Clp_protease_ATP-binding_subunit_ClpA | 0 | 0 | 1 |
|  | fig\|642.770.peg.1312 | NODE_1_length_1964260_cov_90.024368 | Protease_II_(EC_3.4.21.83) | 0 | 0 | 1 |
|  | fig\|642.770.peg.1793 | NODE_1_length_1964260_cov_90.024368 | Uncharacterized_protease_YegQ | 0 | 0 | 1 |
|  | fig\|642.770.peg.1989 | NODE_1_length_1964260_cov_90.024368 | Vibriolysin__extracellular_zinc_protease_(EC_3.4.24.25)_@_Pseudolysin__extracellular_zinc_protease_(EC_3.4.24.26) | 0 | 0 | 1 |
|  | fig\|642.770.peg.2147 | NODE_2_length_947468_cov_91.192050 | Uncharacterized_protease_YdcP | 0 | 0 | 1 |
|  | fig\|642.770.peg.3280 | NODE_3_length_439005_cov_94.027299 | Protease_II_(EC_3.4.21.83) | 0 | 0 | 1 |
|  | fig\|642.770.peg.3795 | NODE_5_length_206066_cov_93.597574 | ATP-dependent_hsl_protease_ATP-binding_subunit_HslU | 0 | 0 | 1 |
|  | fig\|642.770.peg.3796 | NODE_5_length_206066_cov_93.597574 | ATP-dependent_protease_subunit_HslV_(EC_3.4.25.2) | 0 | 0 | 1 |
|  | fig\|642.770.peg.12 | NODE_10_length_72653_cov_95.226534 | Oligopeptidase_A_(EC_3.4.24.70) | 0 | 0 | 1 |
|  | fig\|642.770.peg.67 | NODE_10_length_72653_cov_95.226534 | Xaa-Pro_dipeptidase_PepQ_(EC_3.4.13.9) | 0 | 0 | 1 |
|  | fig\|642.770.peg.308 | NODE_1_length_1964260_cov_90.024368 | Uncharacterized_peptidase_U32_family_member_YhbV | 0 | 0 | 1 |
|  | fig\|642.770.peg.319 | NODE_1_length_1964260_cov_90.024368 | Peptidase_B_(EC_3.4.11.23) | 0 | 0 | 1 |
|  | fig\|642.770.peg.324 | NODE_1_length_1964260_cov_90.024368 | Peptidase_B_(EC_3.4.11.23) | 0 | 0 | 1 |
|  | fig\|642.770.peg.496 | NODE_1_length_1964260_cov_90.024368 | Aminopeptidase_PepA-related_protein | 0 | 0 | 1 |
|  | fig\|642.770.peg.654 | NODE_1_length_1964260_cov_90.024368 | Membrane_alanine_aminopeptidase_N_(EC_3.4.11.2) | 0 | 0 | 1 |
|  | fig\|642.770.peg.1189 | NODE_1_length_1964260_cov_90.024368 | Oligoendopeptidase_F-like_protein | 0 | 0 | 1 |
|  | fig\|642.770.peg.1360 | NODE_1_length_1964260_cov_90.024368 | Tripeptide_aminopeptidase_(EC_3.4.11.4) | 0 | 0 | 1 |
|  | fig\|642.770.peg.1462 | NODE_1_length_1964260_cov_90.024368 | Probable_endopeptidase_NlpC | 0 | 0 | 1 |
| Extracellular DIGESTION of large proteins | fig\|642.770.peg.1502 | NODE_1_length_1964260_cov_90.024368 | FIG009095:_D_D-carboxypeptidase_family_protein | 0 | 0 | 1 |
|  | fig\|642.770.peg.1638 | NODE_1_length_1964260_cov_90.024368 | Peptidase__M23/M37_family | 0 | 0 | 1 |
|  | fig\|642.770.peg.1722 | NODE_1_length_1964260_cov_90.024368 | Membrane_proteins_related_to_metalloendopeptidases | 0 | 0 | 1 |
|  | fig\|642.770.peg.1792 | NODE_1_length_1964260_cov_90.024368 | L_D-transpeptidase_>_YbiS | 0 | 0 | 1 |
|  | fig\|642.770.peg.1981 | NODE_1_length_1964260_cov_90.024368 | L_D-transpeptidase_>_YbiS | 0 | 0 | 1 |
|  | fig\|642.770.peg.2129 | NODE_2_length_947468_cov_91.192050 | Alpha-aspartyl_dipeptidase_Peptidase_E_(EC_3.4.13.21) | 0 | 0 | 1 |
|  | fig\|642.770.peg.2153 | NODE_2_length_947468_cov_91.192050 | Thermostable_carboxypeptidase_1_(EC_3.4.17.19) | 0 | 0 | 1 |
|  | fig\|642.770.peg.2266 | NODE_2_length_947468_cov_91.192050 | Methionine_aminopeptidase_(EC_3.4.11.18) | 0 | 0 | 1 |
|  | fig\|642.770.peg.2291 | NODE_2_length_947468_cov_91.192050 | Peptidase__M13_family | 0 | 0 | 1 |
|  | fig\|642.770.peg.2456 | NODE_2_length_947468_cov_91.192050 | γ-glutamyltranspeptidase_(EC_2.3.2.2)_ @_Glutathione_hydrolase_(EC_3.4.19.13) | 0 | 0 | 1 |
|  | fig\|642.770.peg.3506 | NODE_4_length_411610_cov_92.800192 | Xaa-Pro_aminopeptidase_(EC_3.4.11.9) | 0 | 0 | 1 |
|  | fig\|642.770.peg.3576 | NODE_4_length_411610_cov_92.800192 | Prolyl_endopeptidase_(EC_3.4.21.26) | 0 | 0 | 1 |
|  | fig\|642.770.peg.4017 | NODE_6_length_154685_cov_89.330397 | Oligoendopeptidase_F-like_protein | 0 | 0 | 1 |
|  | fig\|642.770.peg.4147 | NODE_7_length_126480_cov_93.962673 | Bacterial_leucyl_aminopeptidase_(EC_3.4.11.10) | 0 | 0 | 1 |
| DNA DIGESTION | fig\|642.770.peg.760 | NODE_1_length_1964260_cov_90.024368 | Extracellular_and/or_outer_membrane_deoxyribonuclease_NucH/SO1066 | 0 | 0 | 1 |
|  | fig\|642.770.peg.1409 | NODE_1_length_1964260_cov_90.024368 | UPF0294_protein_YafD (exo- and endo- nuclease family) | 0 | 0 | 1 |
|  | fig\|642.770.peg.1884 | NODE_1_length_1964260_cov_90.024368 | DNA/RNA_endonuclease_G | 1 | 1 | 1 |
|  | fig\|642.770.peg.1925 | NODE_1_length_1964260_cov_90.024368 | Extracellular_and/or_outer_membrane_deoxyribonuclease_NucH/SO1066 | 0 | 0 | 1 |
| Peptide TRANSPORT into the cell | fig\|642.770.peg.2135 | NODE_2_length_947468_cov_91.192050 | Succinyl-CoA_synthetase__alpha_subunit | 0 | 0 | 1 |
|  | fig\|642.770.peg.274 | NODE_1_length_1964260_cov_90.024368 | ABC_transporter__permease_protein_1_(cluster_5__nickel/peptides/opines) | 4 | 6 | 3 |
|  | fig\|642.770.peg.275 | NODE_1_length_1964260_cov_90.024368 | ABC_transporter__permease_protein_2_(cluster_5__nickel/peptides/opines) | 3 | 5 | 3 |
|  | fig\|642.770.peg.1213 | NODE_1_length_1964260_cov_90.024368 | Oligopeptide_ABC_transporter__permease_protein_OppC_(TC_3.A.1.5.1) | 4 | 6 | 3 |
|  | fig\|642.770.peg.1214 | NODE_1_length_1964260_cov_90.024368 | Oligopeptide_ABC_transporter__permease_protein_OppB_(TC_3.A.1.5.1) | 4 | 6 | 3 |
|  | fig\|642.770.peg.1215 | NODE_1_length_1964260_cov_90.024368 | Oligopeptide_ABC_transporter__substrate-binding_protein_OppA_(TC_3.A.1.5.1) | 1 | 1 | 1 |
|  | fig\|642.770.peg.1819 | NODE_1_length_1964260_cov_90.024368 | Dipeptide_ABC_transporter__permease_protein_DppC_(TC_3.A.1.5.2) | 4 | 6 | 3 |
|  | fig\|642.770.peg.1820 | NODE_1_length_1964260_cov_90.024368 | ABC_transporter__permease_protein_1_(cluster_5__nickel/peptides/opines) | 4 | 6 | 3 |
|  | fig\|642.770.peg.1913 | NODE_1_length_1964260_cov_90.024368 | ABC_transporter__permease_protein_2_(cluster_5__nickel/peptides/opines) | 4 | 6 | 3 |
|  | fig\|642.770.peg.1914 | NODE_1_length_1964260_cov_90.024368 | ABC_transporter__permease_protein_1_(cluster_5__nickel/peptides/opines) | 4 | 6 | 3 |
| DNA TRANSPORT into the cell | fig\|642.770.peg.2922 | NODE_3_length_439005_cov_94.027299 | Na+_dependent_nucleoside_transporter_NupC | 5 | 9 | 4 |
|  | fig\|642.770.peg.2967 | NODE_3_length_439005_cov_94.027299 | Na+_dependent_nucleoside_transporter_NupC | 4 | 8 | 4 |
|  | fig\|642.770.peg.3112 | NODE_3_length_439005_cov_94.027299 | Na+_dependent_nucleoside_transporter_NupC | 4 | 8 | 4 |
|  | fig\|642.770.peg.4145 | NODE_7_length_126480_cov_93.962673 | Predicted_nucleoside_ABC_transporter__permease_1_component | 6 | 10 | 5 |
|  | fig\|642.770.peg.4146 | NODE_7_length_126480_cov_93.962673 | Predicted_nucleoside_ABC_transporter__permease_2_component | 4 | 7 | 4 |
|  | fig\|642.770.peg.177 | NODE_13_length_22868_cov_91.969299 | DNA_uptake_protein | 0 | 0 | 1 |
